## Supplementary Figures for "ZMYM2 controls human transposable element transcription through distinct co-regulatory complexes"

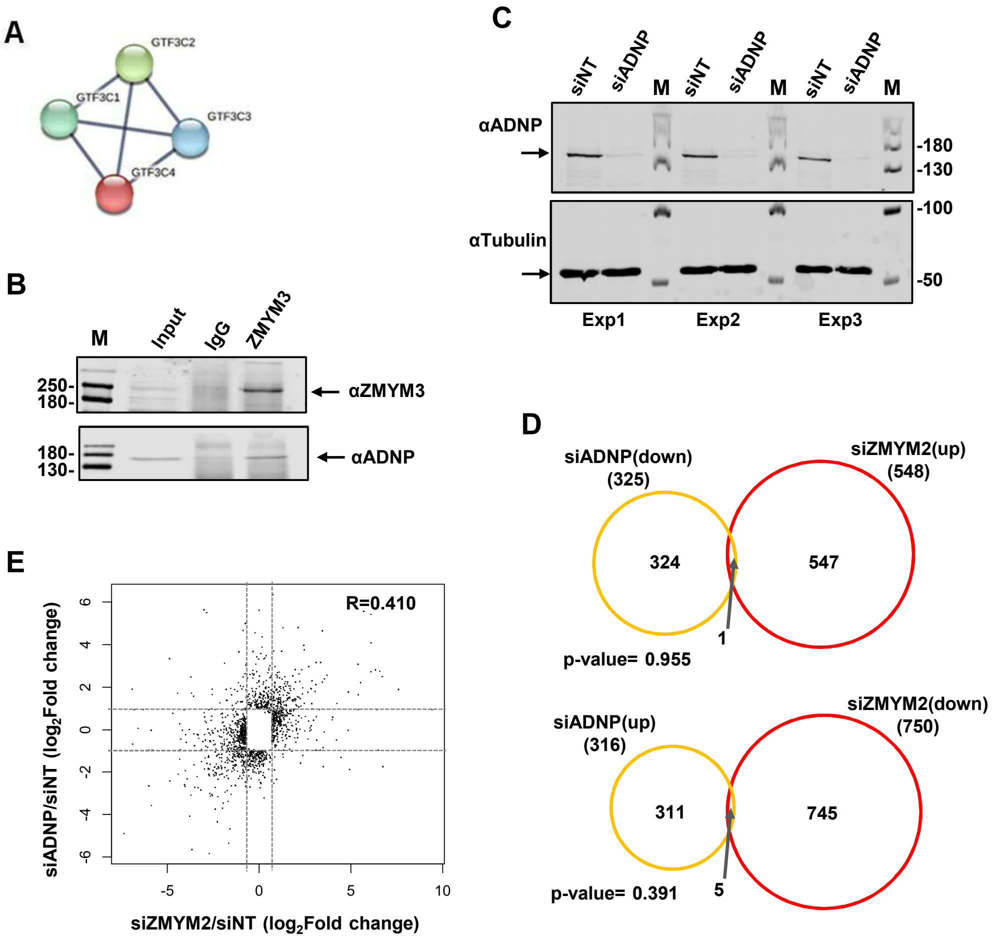


**Fig. S1. ADNP interactions with ZMYM2.** (A) Depiction of interactions between ZMYM2 binding partners found in RIME experiments with known previous protein interactions in the GTF3C complex found in the STRING database (Jensen et al., 2009). (B) Co-immunoprecipitation analysis of ADNP with ZMYM3. Immunoprecipitation (IP) was performed with ZMYM3 or control IgG antibody from U2OS cells and resulting proteins detected by immunoblotting (IB) with the indicated antibodies. Molecular weight markers (M) and 10% input are shown. (C) Western blot of ADNP expression in U2OS cells following treatment with siADNP or a non-targeting (NT) siRNA (top). Tubulin was used as a loading control (bottom). Molecular weight markers (M) are shown. (D) Venn diagrams showing overlaps in genes showing reciprocal directionality following ADNP (left; orange) or ZMYM2 (right; red) depletion. Genes downregulated with siADNP and upregulated with siZMYM2 (top) or upregulated with siADNP and downregulated with siZMYM2 (bottom) are shown (fold change >1.6; Padj <0.01). (E) Scatterplot of significantly changing genes following ZMYM2 (x-axis) or ADNP2 (y-axis) depletion (fold change >1.6; Padj <0.01).

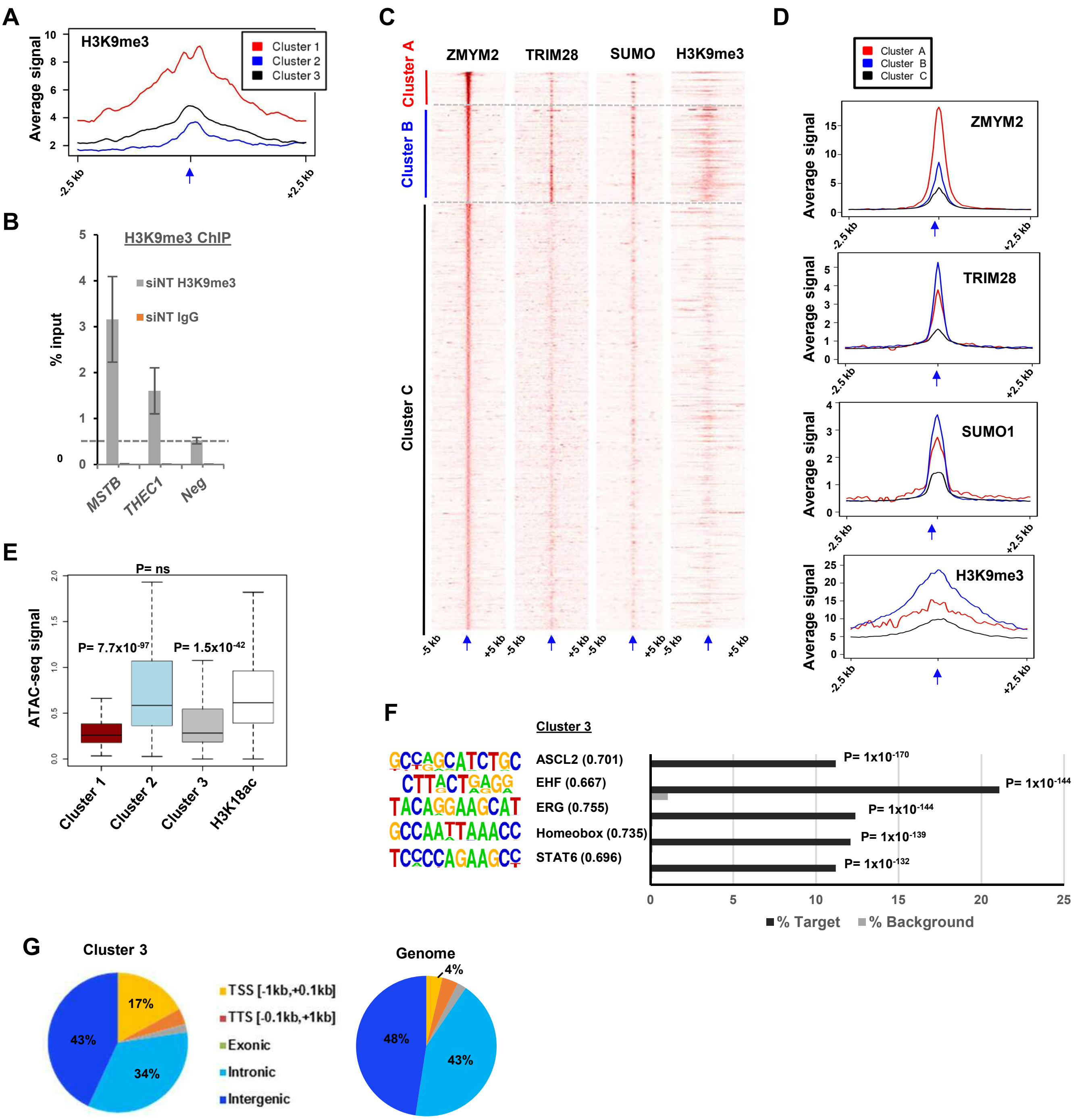


**Fig. S2. Molecular characterisation of ZMYM2 binding regions.** (A) Tag density plot of H3K9me3 ChIP-seq signal from H1 ESCs in the three clusters of ZMYM2 binding regions using ZMYM2 and TRIM28 binding regions for clustering. (B) ChIP-qPCR of H3K9me3 in U2OS cells at the indicated ZMYM2 bound loci or a negative control region (neg) not bound by ZMYM2. Signal from non-specific IgG is also shown at the same regions. Data are shown relative to input (n=3). (C) Heatmaps showing the signals of the indicated proteins or chromatin marks from ChIP-seq experiments in mouse ESCs cells plotted across a 10 kb region surrounding the centres (arrowed) of the ZMYM2 binding regions. Data were clustered to produce 3 clusters. (D) Tag density plots of the indicated ChIP-seq signals in the three clusters of ZMYM2 binding regions. (E) Boxplots showing the ATAC-seq signal at each of the ZMYM2 clusters and regions containing H3K18ac peaks. Significance values are shown relative to the signal at H3K18ac peaks. (F) *De novo* motif analysis of cluster 3 regions. The top five most significantly enriched motifs are shown with motif similarity to the indicated protein shown in brackets. (G) Distribution of binding regions among different genomic categories for cluster 3 and the entire genome.

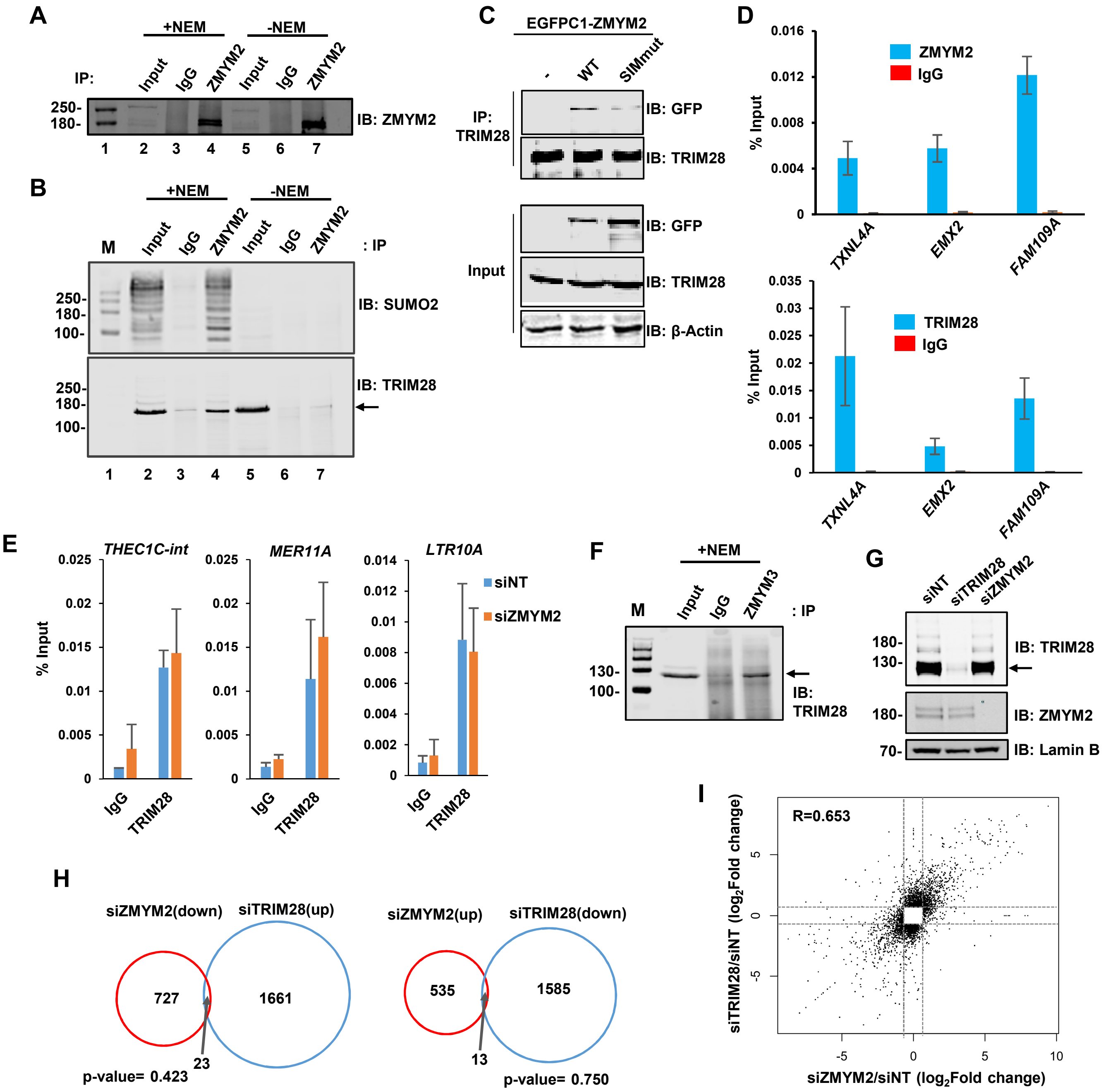


**Fig. S3. ZMYM2-TRIM28 interactions.** (A) Co-immunoprecipitation analysis of TRIM28 with ZMYM2. This is a longer exposure of Fig. 4A top panel to better visualize the input protein. (B) Co-immunoprecipitation analysis of TRIM28 or SUMO2 with ZMYM2. Immunoprecipitation (IP) was performed with ZMYM2 or control IgG antibody from U2OS cells and the resulting proteins detected by immunoblotting (IB) with the indicated antibodies. 10% input is shown. NEM was added to the extracts where indicated. (C) Co-immunoprecipitation (IP) analysis of endogenous TRIM28 with the indicated EGFP-tagged ZMYM2 proteins. Inputs (bottom) and IPs (top) were immunoblotted (IB) with the indicated antibodies. (D) ChIP-qPCR of ZMYM2 (top) and TRIM28 (bottom) in U2OS cells at the indicated ZMYM2 bound loci. Signal from non-specific IgG is also shown at the same regions. Data are shown relative to input (n=3). (E) ChIP-qPCR of TRIM28 in U2OS cells following treatment with siZMYM2 or a non-targeting (NT) siRNA, at the indicated ZMYM2 bound loci. Signal from non-specific IgG is also shown at the same regions. Data are shown relative to input (n=2). (F) Co-immunoprecipitation analysis of TRIM28 with ZMYM3. Immunoprecipitation (IP) was performed with ZMYM2 or control IgG antibody from U2OS cells in the presence of NEM and the resulting TRIM28 detected by immunoblotting (IB). 10% input is shown. (G) Western blot analysis of lysates from U2OS cells treated with the indicated targeting or non-targeting (NT) siRNAs. Lamin B (loading control), TRIM28 and ZMYM2 were detected by immunoblotting (IB). (H) Venn diagrams showing overlaps in genes showing reciprocal directionality following ZMYM2 (left circle) or TRIM28 (right circle) depletion. Genes downregulated with siTRIM28 and upregulated with siZMYM2 (left) or upregulated with siTRIM28 and downregulated with siZMYM2 (right) are shown (fold change >1.6; Padj <0.01). (I) Scatterplot of significantly changing genes following ZMYM2 (x-axis) or TRIM28 (y-axis) depletion (fold change >1.6; Padj <0.01).

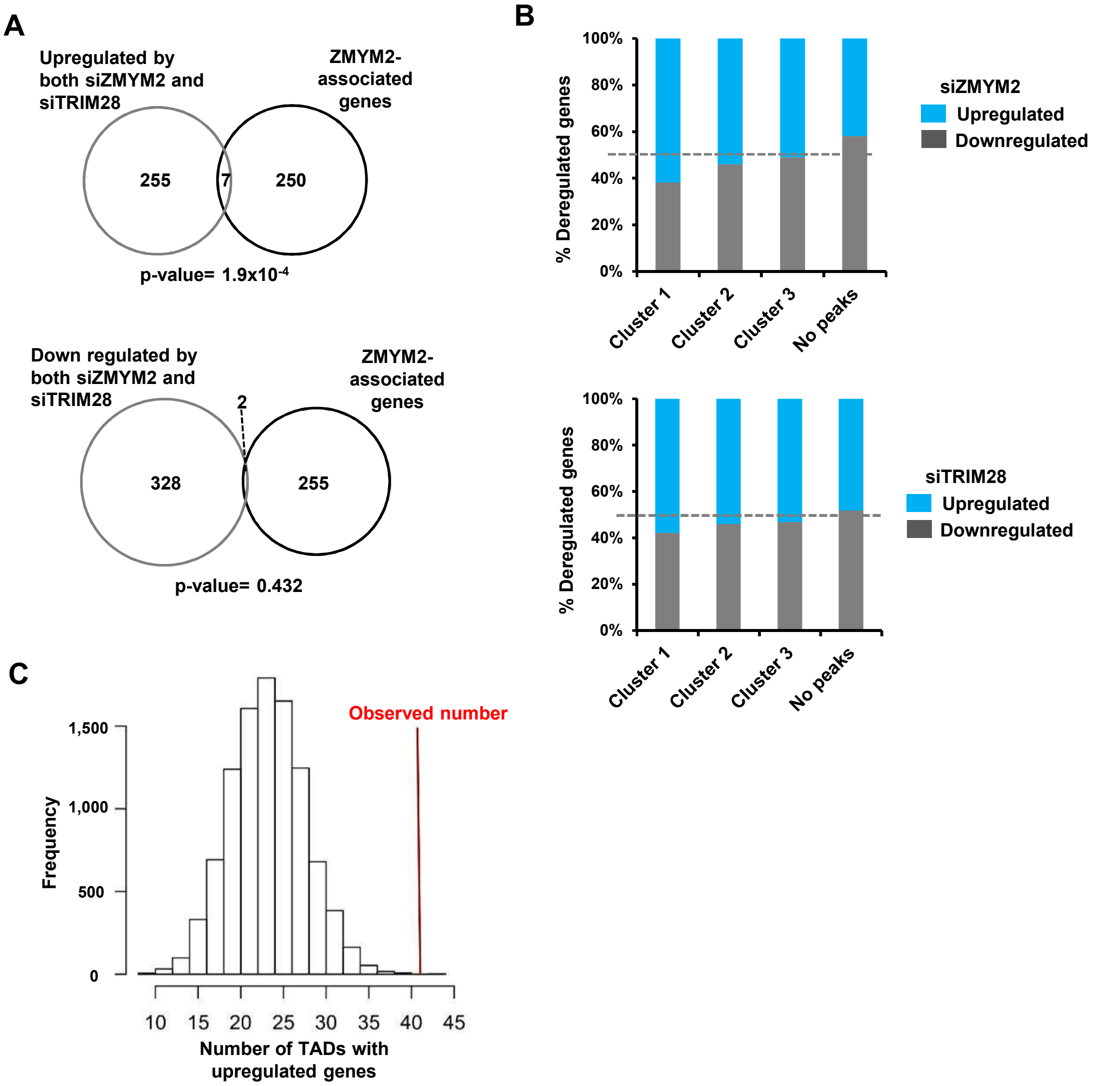


**Fig. S4. ZMYM2 location and gene expression.** (A) Overlap between the closest genes to ZMYM2 binding regions from cluster 1 and the intersection of differentially expressed genes following ZMYM2 or TRIM28 depletion. (B) Relative proportion of genes up- or down-regulated following either ZMYM2 (top) or TRIM28 (bottom) depletion in TADs which also contain ZMYM2 peaks from the indicated clusters or have no ZMYM2 peaks in the same TADs. (C) Frequency distribution of the number of TADs containing a gene upregulated (fold change >1.6; Padj <0.01) following ZMYM2 depletion. 10,000 iterations were performed by randomly selecting 216 TADs across all 3062 TADs. The observed number of TADs containing an upregulated gene (42) from the 216 TADs containing a cluster 1 ZMYM2 peak is shown (P-value = 0.0002).

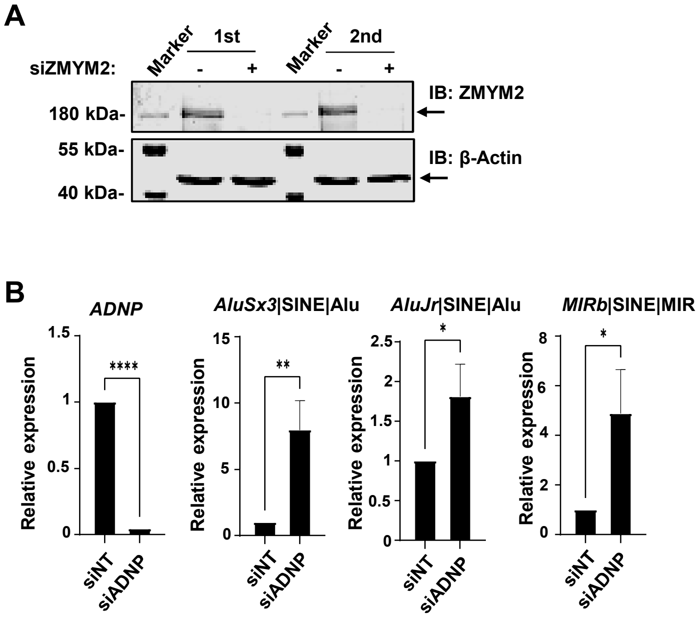


**Fig. S5. ZMYM2 containing complexes regulate retrotransposons.** (A) Western blot illustrating the knockdown efficiencies of ZMYM2 in the RT-qPCR experiments. β-actin is shown as a loading control and sizes of molecular weight markers are indicated. (B) RT-qPCR analysis of expression of ADNP and the indicated SINE elements following ADNP depletion or control non-targeting (NT) siRNA treatment. Individual paired experiments are shown (n=3; P-values *= <0.05, **=<0.01, ****=<0.0001).
