## Supplementary Table1 for "ZMYM2 controls human transposable element transcription through distinct co-regulatory complexes"

| PRIMARY ANTIBODIES | SUPPLIER | SPECIES | TYPE | APPLICATION | REFERENCE |
| --- | --- | --- | --- | --- | --- |
| ZMYM2 (ZNF198) | Bethyl Laboratories | Rabbit | Polyclonal | WB, IP | A301-711A |
| ZMYM3 | Abcam | Rabbit | Polyclonal | WB, IP | Ab106626 |
| ADNP | Abcam | Rabbit | Polyclonal | WB, IP, ChIP-seq | Ab231950 |
| TRIM28  (KAP1) | Abcam | Rabbit | Polyclonal | WB, PLA,  ChIP-seq | Ab10483 |
| SUMO2/3 | Abcam | Rabbit | Polyclonal | WB | Ab3742 |
| MYC | Santa Cruz Biotechnology | Mouse | Monoclonal | PLA | SC-40 |
| H3K9me3 | Abcam | Rabbit | Polyclonal | ChIP | Ab8898 |
| TUBLIN | Sigma-Aldrich | Mouse | Monoclonal | WB | T9026-2ML |
| LAMIN B | Santa Cruz Biotechnology | Goat | Polyclonal | WB | SC-6216 |
| β-ACTIN | Millipore | Mouse | Monoclonal | WB | MAB1501 |
| IgG | Millipore | Rabbit | - | IP, ChIP | 12-370 |
| GFP | Santa Cruz Biotechnology | Mouse | Polyclonal | WB | sc-8334 |

| SECONDARY ANTIBODIES | SUPPLIER | SPECIES | TYPE | APPLICATION | REFERENCE |
| --- | --- | --- | --- | --- | --- |
| **IRDye 800CW Anti Rabbit** | LI-COR Biosciences | Donkey | Polyclonal | WB | 926-32213 |
| **IRDye 800CW Anti Mouse** | LI-COR Biosciences | Goat | Polyclonal | WB | 926-32210 |
| **IRDye 680LT Anti-Goat** | LI-COR Biosciences | Donkey | Polyclonal | WB | 925-68024 |

**Supplementary Table S1**. List of antibodies
